# Design of the recombinant influenza neuraminidase antigen is crucial for protective efficacy

**DOI:** 10.1101/2021.04.29.442077

**Authors:** Jin Gao, Laura Klenow, Lisa Parsons, Tahir Malik, Je-Nie Phue, Zhizeng Gao, Stephen G. Withers, John Cipollo, Robert Daniels, Hongquan Wan

## Abstract

Supplementing influenza vaccines with recombinant neuraminidase (rNA) remains a promising approach for improving the suboptimal efficacy. However, correlations among rNA designs, properties, and protection have not been systematically investigated. Here, we performed a comparative analysis of several rNAs produced from different construct designs using the baculovirus/insect cell system. The rNAs were designed with different tetramerization motifs and NA domains from a recent H1N1 vaccine strain (A/Brisbane/02/2018) and were analyzed for enzymatic properties, antigenicity, thermal and size stability, and protection in mice. We found that rNAs containing the NA head-domain versus the full-ectodomain possess distinct enzymatic properties and that the molecular size stability is tetramerization domain-dependent, whereas protection is more contingent on the combination of the tetramerization and NA domains. Following single-dose immunizations, a rNA possessing the full-ectodomain, non-native enzymatic activity, and the tetramerization motif from the human vasodilator-stimulated phosphoprotein provided substantially higher protection than a rNA possessing the head-domain, native activity and the same tetramerization motif. In contrast, these two rNAs provided comparable protection when the tetramerization motif was exchanged with the one from the tetrabrachion protein. These findings demonstrate that the rNA design is crucial for the protective efficacy and should be thoroughly evaluated for vaccine development, as the unpredictable nature of the heterologous domain combination can result in rNAs with similar key attributes but vastly differ in protection.

**IMPORTANCE:** For several decades it has been proposed that influenza vaccines could be supplemented with recombinant neuraminidase (rNA) to improve the efficacy. However, some key questions for manufacturing stable and immunogenic rNA remain to be answered. We show here that the tetramerization motifs and NA domains included in the rNA construct design can have a profound impact on the biochemical, immunological and protective properties. We also show that the single-dose immunization regimen is more informative for assessing the rNA immune response and protective efficacy, which is surprisingly more dependent on the specific combination of NA and tetramerization domains than common attributes for evaluating NA. Our findings may help to optimize the design of rNAs that can be used to improve or develop influenza vaccines.

## INTRODUCTION

Influenza vaccine efficacy remains suboptimal despite the concerted efforts to monitor the evolution of influenza viruses and frequent updates on the vaccine composition (1–3). A contributing factor to the poor efficacy is the need to prepare candidate vaccine viruses containing a hemagglutinin (HA) that antigenically matches the circulating strains months ahead of each influenza season. This requirement combined with the propensity of HA to mutate under selective pressure from antibodies makes it difficult to significantly improve the vaccine efficacy using HA alone. Accordingly, many studies have begun to investigate the potential benefits of including other influenza antigens such as neuraminidase (NA), the second most abundant surface glycoprotein on influenza virus (4, 5).

NA is a sialidase that promotes the spread of influenza virus by removing the receptors for HA (5, 6). Previous work has shown that NA-specific antibodies/immunity can inhibit the growth of influenza viruses *in vitro* and confer protection against influenza virus infection in animal models and humans (7–12). The confirmed benefits of NA immunity and additional advantages such as its relatively slower evolution than HA (13, 14), make NA an attractive target for optimizing influenza vaccines.

Current inactivated influenza vaccines often contain the NA from the recommended vaccine strains. However, the amount is usually low and variable (15, 16), likely due to its labile nature and strain-dependent differences in NA content (17). Options for addressing this bottleneck include developing candidate vaccine viruses that contain higher NA content or supplementing influenza vaccines with purified viral NA or recombinant NA (rNA). While NA isolated from viruses and produced recombinantly have both shown promising protective efficacy (18–24), rNA expressed in the baculovirus/insect cell system currently has a greater potential for practical use because of its capacity to generate high yields and the system is currently used for manufacturing licensed vaccines, including the HA-based influenza vaccine, Flublok (25).

Prior studies examining rNA protection have tested various construct designs and have all used a two-dose immunization (prime and boost) with rNA protein amounts as high as >20 μg per mouse (26–30). While these studies have demonstrated the protective benefit of rNA, several key questions remain for implementing rNA antigens in influenza vaccines, *e.g.*, How does the rNA construct design affect the quality attributes and protective efficacy? Is NA enzymatic activity a reliable indicator of rNA immunogenicity/protection? Can protection be achieved with a single dose of rNA using cost-effective amounts? In the current study we have addressed these questions using rNAs that contain different stabilizing tetramerization domains combined with either the NA head-domain or full-ectodomain. Our results show that the rNA construct design is critical for protection by single-dose immunization with low rNA amounts and that rNA antigens being developed for influenza vaccines should be thoroughly characterized, as the protective efficacy can differ between rNAs with similar quality attributes.

## RESULTS

### Design and purification of rNAs expressed in insect cells

Influenza NA is a membrane glycoprotein that functions on the viral surface as a homotetramer (Fig. 1A) (31, 32). It is comprised of an enzymatic head-domain connected to a short stalk region and an *N*-terminal transmembrane domain. Due to the low abundance of NA in virions, various designs and approaches have been used to generate rNA for structural and immunological studies as well as vaccine development (21, 33–36). To examine if the rNA construct design correlates with the biochemical properties and protective efficacy, we expressed four secreted, soluble rNAs using the NA sequence from an H1N1 vaccine strain, A/Brisbane/02/2018 (N1-BR18). The constructs were designed based on a common approach that includes the addition of a signal peptide, a 6×His-tag, and a tetramerization motif in place of the *N*-terminal transmembrane domain of NA (Fig. 1B) (33, 34). For two of the constructs, we combined the tetramerization domain from the human vasodilator-stimulated phosphoprotein (VASP) with either the full-ectodomain of N1-BR18 (V35), or the head-domain (V82), and the remaining two (T35 and T82) followed a similar design using the tetrabrachion (TB) tetramerization domain instead (Fig. 1B).

**Figure 1.**
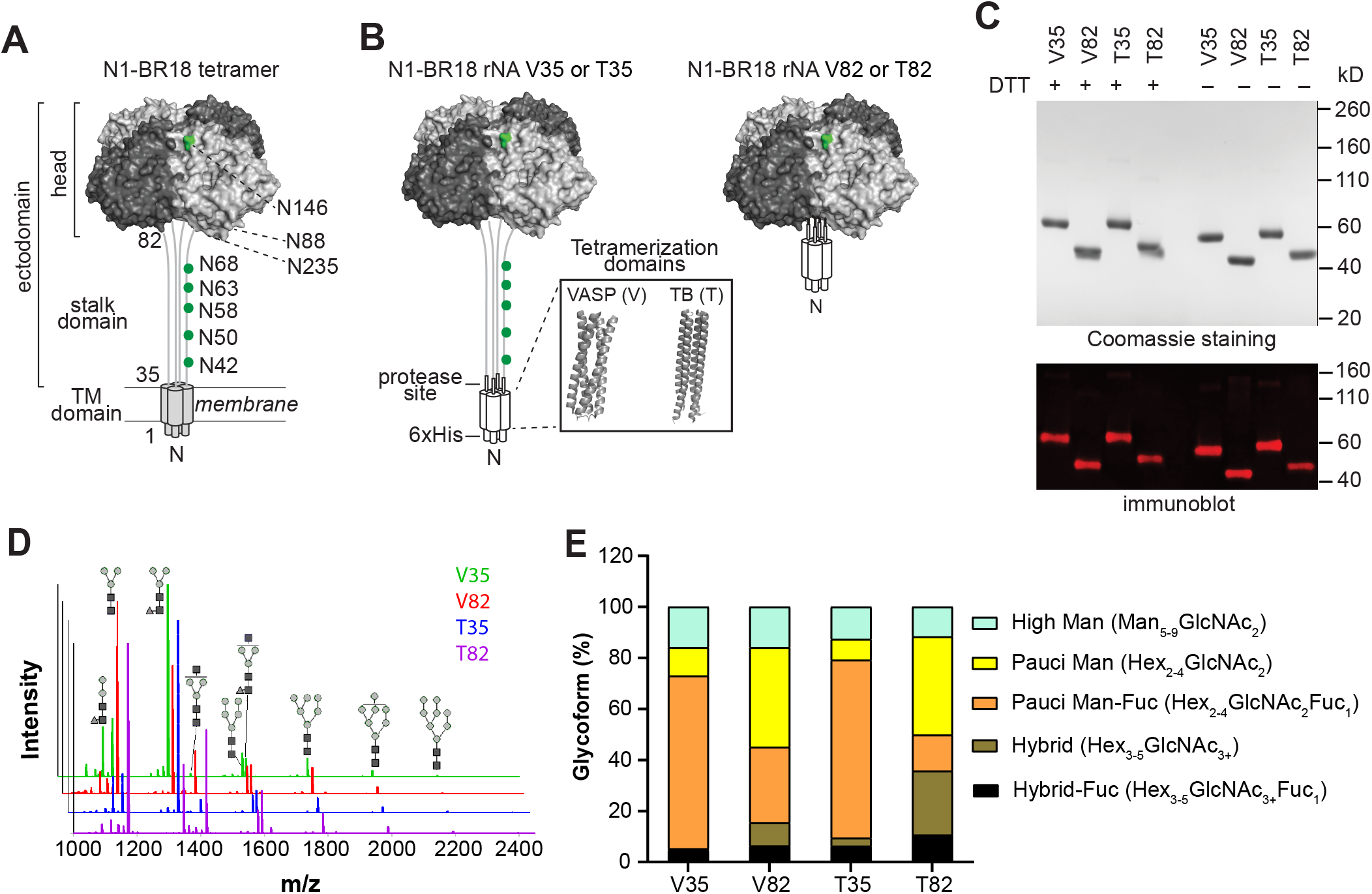
Characterization of N1-BR18 rNAs. **(A)** A schematic diagram of full-length N1-BR18 showing the trans-membrane (TM) domain and the ectodomain, which includes the NA head and stalk. Potential *N*-linked glycosylation sites are labeled (green). Sites 88 and 235 are not visible in the displayed view. **(B)** Diagrams of the N1-BR18 rNA construct designs. V35 and T35 contain the full-ectodomain of N1-BR18 (residues 35-469) connected to the tetramerization domains from VASP (V35) or TB (T35). V82 and T82 are designed similarly using the N1-BR18 head-domain (residues 82-469). Structures of tetramerization domains from VASP (PDB ID: 1USE) and TB (PDB ID: 1FE6) are shown in a box. **(C)** Representative images of a Coomassie stained SDS-PAGE gel containing the rNAs (2 μg/lane) and an immunoblot (0.2 μg NA/lane) resolved using a N1-specific mAb. The rNAs were untreated or reduced with DTT prior to resolution by SDS-PAGE. **(D)** Spectra of PNGase F-released *N*-linked permethylated glycans from each rNA. Structures of the most abundant glycoforms are shown, mannose (grey circles), *N*-acetyl glucosamine (black squares) and fucose (grey triangle). **(E)** Graph displaying the abundance of the different glycoform subtypes. Mannose, Man; hexose, Hex; *N*-acetyl glucosamine, GlcNAc; fucose, Fuc.

The rNAs were expressed using High Five insect cells and isolated from the culture medium by immobilized metal affinity chromatography. Following the isolation, the four rNAs resolved at the expected molecular weight by SDS-PAGE, showed high purity based on Coomassie staining, and reacted with an N1-specific monoclonal antibody (mAb) by immunoblotting (Fig. 1C). In the absence of reductant (dithiothreitol, DTT), each rNA displayed faster mobility on SDS-PAGE, suggesting that they possess the proper intramolecular disulfide bonds.

Since *N*-linked glycans can influence antigenicity and NA folding (4, 37, 38), we also analyzed the *N*-linked glycoforms on the rNAs by mass-spectrometry. As expected for a glycoprotein produced by insect cells, the majority of the *N*-linked glycans on the rNAs were small and mainly consisted of pauci-mannose and high mannose glycoforms (Fig. 1D). Both rNAs (V82 and T82) containing the head-domain showed a higher abundance of pauci-mannose and complex glycoforms (Fig. 1E), whereas fucosylated pauci-mannose glycoforms were more prevalent on the rNAs (V35 and T35) comprised of the full-ectodomain (Fig. 1E), suggesting these are from the stalk region. Interestingly, the glycoform distribution somewhat differed between V82 and T82, but not V35 and T35, indicating that the tetramerization domain may influence glycosylation of the smaller rNA constructs.

### Enzymatic properties of N1-BR18 rNAs

NA only functions in its native tetrameric conformation, suggesting sialidase activity is a reasonable indicator for the proper NA structural conformation (32). To compare the sialidase activity of the rNAs we used the synthetic substrate 2’-(4-methylumbelliferyl) α-D-N-acetylneuraminic acid (Mu-NANA). The rNAs with the head-domain (V82 and T82) showed an activity that was ~10-fold higher than the rNAs with the full-ectodomain (V35 and T35) and much closer to that observed for a similar amount of full-length N1-BR18 in purified reassortant virus (BR18×WSN) that bears the HA and NA genes from BR18 and the internal genes from the H1N1 strain A/WSN/1933 (Fig. 2A). A Michaelis-Menten kinetic analysis revealed that the lower activity of V35 and T35 is not associated with a change in the substrate binding affinity, as the *K*_m_ values of the four rNAs were similar (Fig. 2B and Table 1).

**TABLE 1.** Enzymatic properties of N1-BR18 rNAs.

| rNAs | $K_m$<br>($\mu\text{M}$) | Apparent $k_{\text{cat}}$<br>($\text{s}^{-1}$) | Functional rNAs<br>(%) | True $k_{\text{cat}}$<br>($\text{s}^{-1}$) |
| --- | --- | --- | --- | --- |
| V35 | 48.6<sup>a</sup> | 12 | 6.8 | 176 |
| V82 | 43.1 | 130 | 50 | 260 |
| T35 | 45.6 | 10 | 4.3 | 232 |
| T82 | 42.7 | 123 | 46 | 267 |
| Virus<sup>b</sup> | 44.6 | 221 | 74 | 298 |

<sup>a</sup>Mean from three technical repeats.
<sup>b</sup>Purified BR18×WSN virus, which contains ~10% NA, was used as a control.

^*a*^Mean from three technical repeats. ^*b*^Purified BR18×WSN virus, which contains ~10% NA, was used as a control.

**Figure 2.**
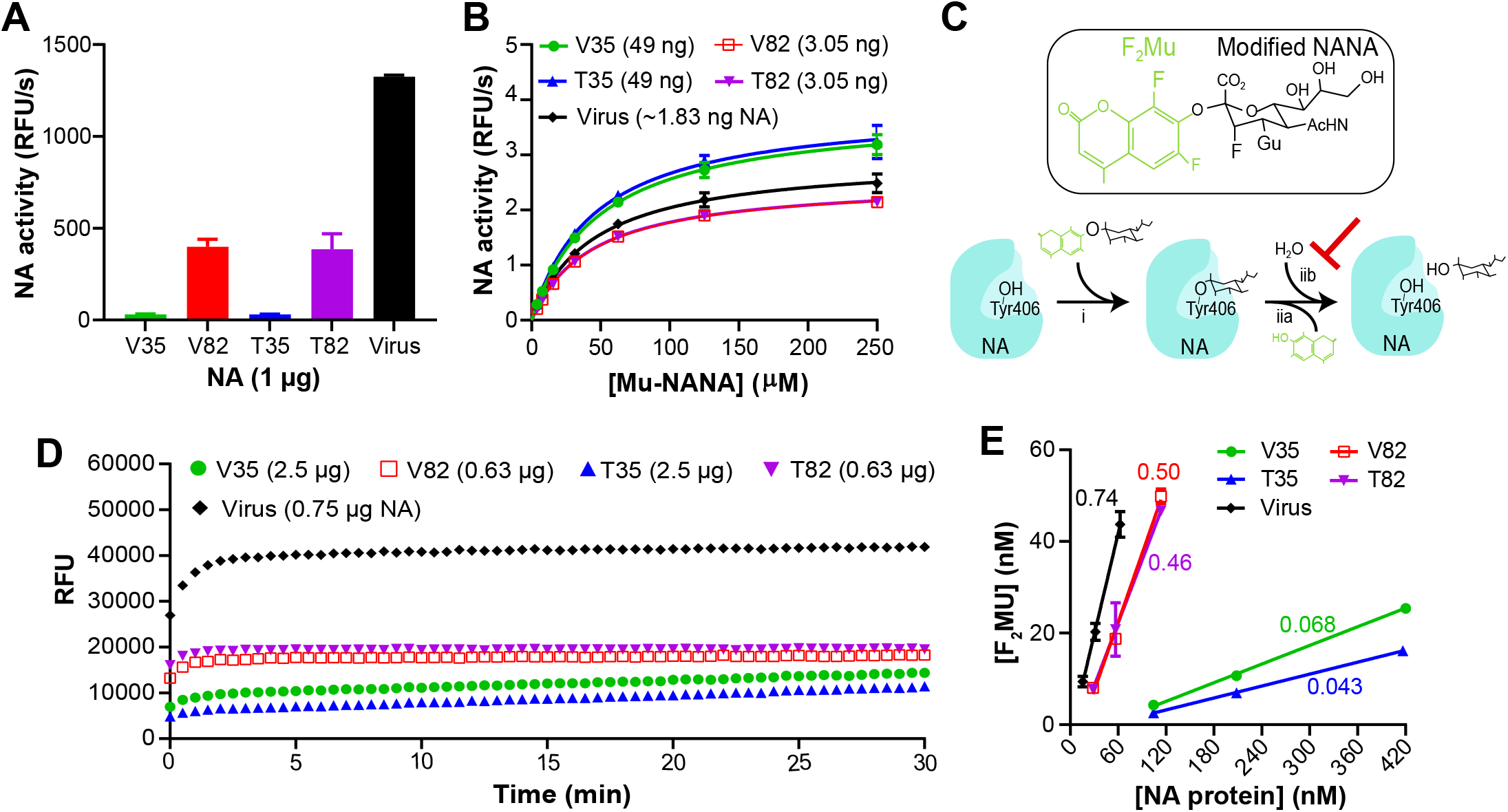
Designs with the N1-BR18 head-domain produce more active rNA. **(A)** Graph displaying the mean enzymatic activity of the indicated rNAs that was determined using the synthetic substrate Mu-NANA. The activity is expressed as relative fluorescence unit per sec (RFU/s) and corresponds to the value from 1 μg of rNA. Purified BR18×WSN virus (a reassortant carrying the HA and NA genes from BR18 and the internal genes from H1N1 A/WSN/1933) containing an equivalent amount of NA was included for comparison. Error bars represent the standard deviation (SD) from three technical repeats. **(B)** Michaelis-Menten kinetic analysis of N1-BR18 rNAs. The activity of the indicated amounts of the rNAs was measured using increasing concentrations of Mu-NANA. The mean value is shown ± SD from three technical repeats. **(C)** Diagram showing the reaction of TR1 (structure in upper panel) with NA. The NA catalytic residue (Tyr406) makes a nucleophilic attack of the TR1 reagent (i), resulting in the release of F2Mu and a covalently bound TR1 sialic acid intermediate (iia). The presence of the guanidinium (Gu) and fluorine modifications decrease the subsequent attack by H2O (iib), which facilitates the sialic acid release. **(D)** Graph displaying the mean fluorescent measurements from the reaction of the indicated rNA amounts with the TR1 reagent from three independent runs using 30 sec intervals. **(E)** Correlation plot showing the protein concentrations of each rNA and the F2Mu concentration that was released from the TR1 reagent. The linear regression slopes, used to determine the fraction of active rNA in each preparation, are displayed. The data are means ± SD from three technical repeats. Purified BR18×WSN virus was included in all assays for comparison.

To determine whether V35 and T35 possess lower catalytic rates (*k*_cat_) or a smaller percentage of enzymatically active rNA in the preparations, we analyzed the rNA preparations using the active-site titrating agent TR1 (39, 40). TR1 is a modified Mu-NANA compound that only undergoes a single sialic acid cleavage reaction per enzyme molecule (Fig. 2C), releasing one equivalent of difluoromethylumbelliferyl alcohol (F_2_Mu) in a burst phase, which can be followed by a slow steady-state turnover phase for some NAs (40). The TR1 profiles for V82 and T82 resembled full-length N1-BR18 with a high initial burst that reached a maximum within ~2 minutes (Fig. 2D). V35 and T35 at higher protein amounts both showed a smaller initial burst followed by a slow steady-state turnover phase, indicating these rNAs possess a lower proportion of enzymatically active tetramers that likely possess an alteration in the active site, which can facilitate release of the covalently bound TR1 intermediate. We then calculated the fraction of enzymatically active rNA in each preparation by plotting the concentration of F_2_Mu released from TR1 by each rNA at three protein concentrations (Fig. 2E). The results showed that ~50% (slope = ~0.5) of the V82 and T82 preparations are enzymatically active, in line with the percentage (~74%) observed for full-length N1-BR18. In contrast, only ~5% (slope = ~0.05) of the V35 and T35 preparations reacted with TR1 (Fig. 2E and Table 1). Using these values to derive the concentrations of active enzyme, the calculated *k*_cat_ was found to be similar for all four rNAs (Table 1). These results confirm that the designs for T35 and V35 produce ~90% less functional NA than the V82 and T82 designs, indicating that the proportion of functional rNA is mainly influenced by the NA domain rather than the tetramerization domain.

### Analysis of head-domain epitopes on N1-BR18 rNAs

Based on the significant differences in the amount of functional NA, we analyzed the antigenic integrity of the rNAs by a sandwich ELISA (41) using mAbs CD6, 4C4, 1H5 and 4E9, which bind various regions (42–44) in the N1 head-domain (Fig. 3A). The rNAs were first bound with CD6, 4C4 or 1H5 and then detected using the HRP-conjugated mAbs 4C4 or 4E9. At an arbitrary concentration of 1 μg/ml all four rNAs were readily bound by these mAbs and the signals were in line with those obtained for an equivalent amount of full-length N1-BR18 in purified virions (Fig. 3B). To confirm if these epitopes are preserved over time, rNAs stored at 4°C for ~4 months were serially diluted and tested in ELISA with mAbs CD6 and 4E9. All the rNAs were effectively detected at concentrations as low as 7.8-62.5 ng/ml (0.78-6.25 ng/well). V35 and V82 were even detected at lower concentrations (Fig. 3C), suggesting the head-domain epitopes are slightly better conserved with the VASP tetramerization domain, or that these rNAs possess a different property than T35 and T82. Despite the subtle differences, the overall similarity of the binding profiles implies that the head-domain epitopes remain largely intact on the rNAs even though the percentage of functional rNA differ.

**Figure 3.**
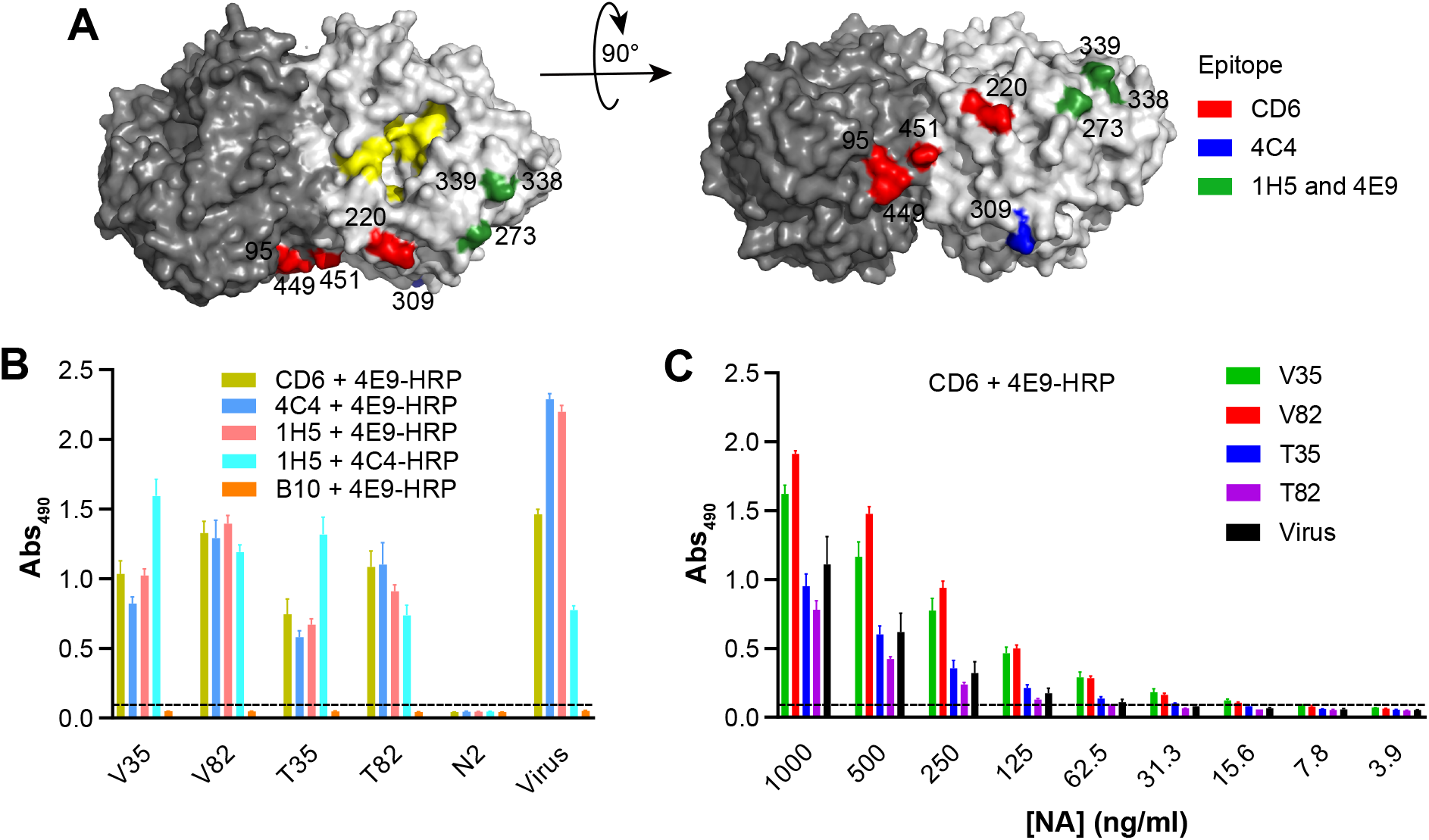
N1-BR18 rNAs retain the antigenicity of multiple head domain epitopes. **(A)** Side (left panel) and top views (right panel) of an N1 dimer (PDB ID: 3NSS) showing the three epitopes that are recognized by the mAbs CD6 (red), 4C4 (blue), and the broadly reactive mAbs 1H5 and 4E9 (green). The NA active site residues 118, 151, 152, 224, 276, 292, 371 and 406 are shown in yellow. **(B)** N1-BR18 rNAs were readily bound by mAbs CD6, 4C4, 1H5, and 4E9. Binding was measured by a sandwich ELISA, in which mAbs CD6, 4C4, and 1H5 were used to capture the rNAs and the HRP-conjugated mAbs 4E9 (4E9-HRP) and 4C4 (4C4-HRP) were used for detection. An N2-specific mAb B10 and a rNA from the strain A/Minnesota/11/2010 (H3N2) were used as negative controls. rNAs were tested at 1 μg/ml. The data are means ± SD from three technical repeats. **(C)** Serially diluted N1-BR18 rNAs were captured with mAb CD6 and detected with the 4E9-HRP. The data are means ± SD from three technical repeats. Purified BR18×WSN virus was included as a control

### Stability and molecular size analysis of N1-BR18 rNAs

Stability is a crucial attribute for vaccine antigens and NA is known to be a labile tetrameric enzyme (17). To test for stability differences, each rNA was examined by a thermal denaturation analysis and after freeze-thaw-cycling using enzymatic activity as a read-out. All the rNAs showed similar thermostability profiles and the *T_50_* (temperature at which the enzymatic activity was reduced by 50%) values were ~57°C (Fig. 4A), far above routine vaccine manufacturing and storage temperatures. Following multiple freeze-thaw cycles, the rNAs also did not show evident activity loss (Fig. 4B), indicating the stability of the functional rNAs is similar for each design.

**Figure 4.**
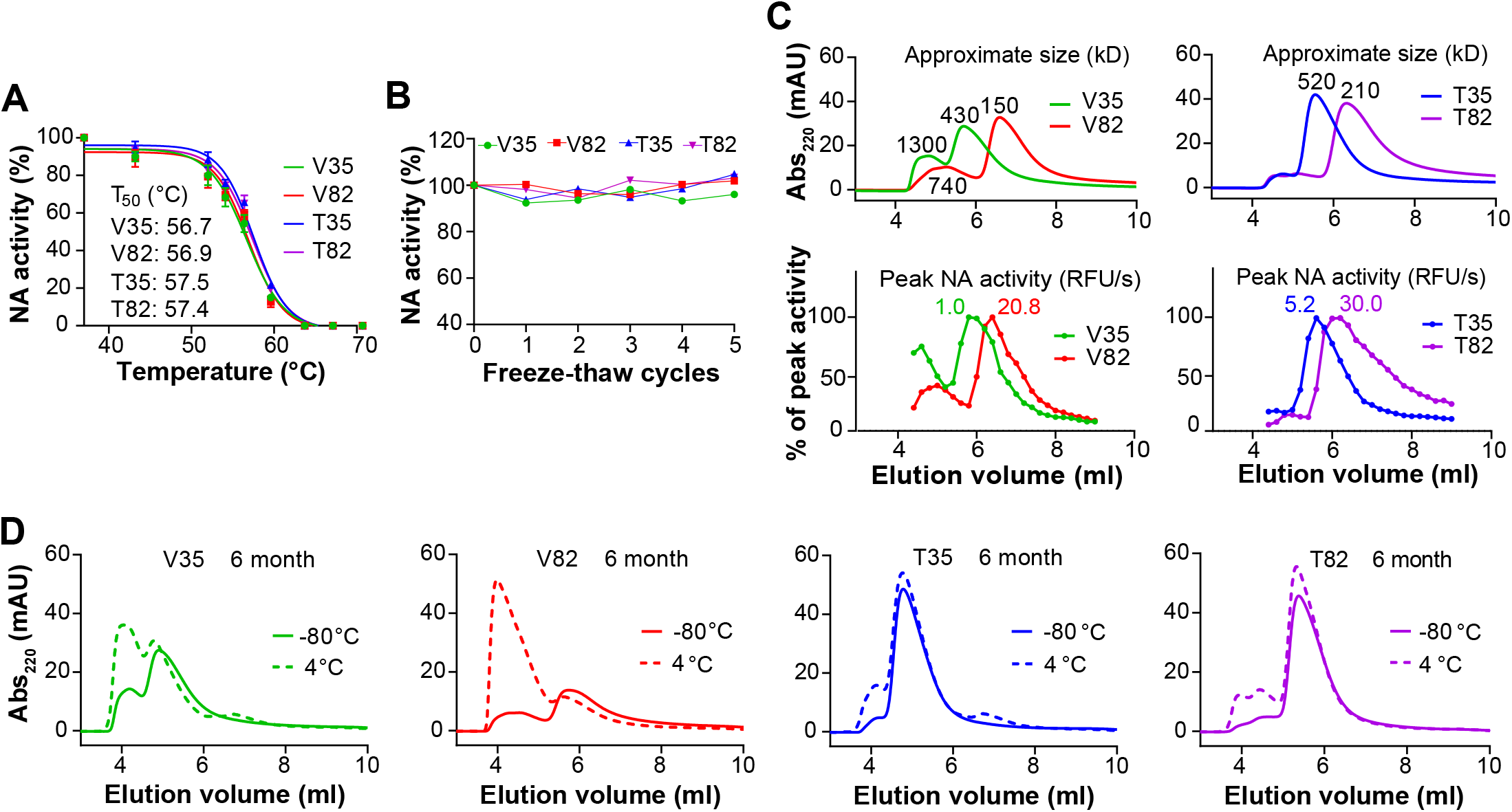
Stability similarities and variations between N1-BR18 rNAs. **(A)** Graphs displaying the thermal melt curves for the rNAs. The rNAs were incubated for 10 min at the indicated temperatures, the enzymatic activity was measured and normalized using the activity of the sample at 37°C. Shown are the mean ± SD of the data from three technical repeats together with the *T_50_* value for each rNA. **(B)** The rNAs were subjected to 5 cycles of freezing on dry ice for 10 min and thawing briefly at 37°C. The enzymatic activity was measured following each freeze-thaw and normalized using the activity from an untreated sample. Means from three technical repeats are displayed. **(C)** SEC profiles of the rNAs shortly after purification. The rNAs were adjusted to 0.5 mg/ml and equal volumes were loaded onto an SEC column. The absorbance at 220 nm (Abs220) versus the elution volume is shown with the estimated molecular weights corresponding to each peak (upper panels). The NA activity profiles of fractions collected between 4.0 and 8.8 ml are shown in the bottom panels, with the activity of each fraction being normalized to the peak activity, which is displayed for each rNA. **(D)** SEC profiles of the rNAs stored at −80°C or 4°C for 6 months.

The molecular size of the rNAs was monitored by size-exclusion chromatography (SEC) within 10 days post-purification and after storage at −80°C and 4°C for 1, 3 and 6 months. In agreement with the estimated molecular weights, the newly purified V82 was found to have the smallest molecular size followed by T82, V35 and T35 (Fig. 4C, upper panel). V35 and V82 also showed a more prominent early peak, corresponding to a larger molecular size, which tracked with the NA activity readings (Fig. 4C, lower panel). While all the SEC profiles were similar to the newly purified rNAs following long-term storage at −80°C, only T35 and T82 showed clear size stability after storage at 4°C for 6 months (Fig. 4D). In contrast, V35 and V82 exhibited dramatic shifts in the SEC profiles at 4°C and the shifts became more prominent in a time-dependent manner (Fig. 4D), indicating that the VASP tetramerization motif promotes the formation of higher order oligomers or multimers, likely explaining the higher sensitivity of V35 and V82 observed in ELISA (Fig. 3C). Despite the extensive molecular size shifts, no loss in enzymatic activity was observed for V35 or V82 after storage at 4°C (data not shown). These findings demonstrate that the rNAs with the TB tetramerization motif possess a more stable molecular size than those with the VASP tetramerization motif.

### Evaluation of the antibody response and protection elicited by the N1-BR18 rNAs

The immunogenicity and protective efficacy of rNAs have been evaluated using a two-dose (prime and boost) approach in animal models (26, 28, 45, 46). To assess the impact of the construct design on the rNA protective efficacy, we immunized mice with the rNAs and conducted lethal viral challenge (Fig. 5A). For the initial evaluation, we measured the NA antibody response in mice that received either two intramuscular (i.m.) immunizations with 5 μg of rNA adjuvanted with poly(I:C) or one i.m. immunization with 5 μg or 2 μg of rNA adjuvanted with poly(I:C). Based on the ELISA results using full length N1-BR18 in purified virus as an antigen, the serum NA-binding antibody titers were higher in mice that received two immunizations (Fig. 5B). Mice that were immunized with a single dose showed consistent differences in the NA-binding antibody titers from each rNA with V35 eliciting the strongest response, followed by T82, T35 and V82 (Fig. 5B). The serum NA-inhibition (NAI) antibody titers, which are considered indicative of protection (12, 16), displayed a similar pattern where V35 elicited the highest NAI titers, and V82 elicited the lowest with all NAI titers below the limit of detection (Fig. 5C).

**Figure 5.**
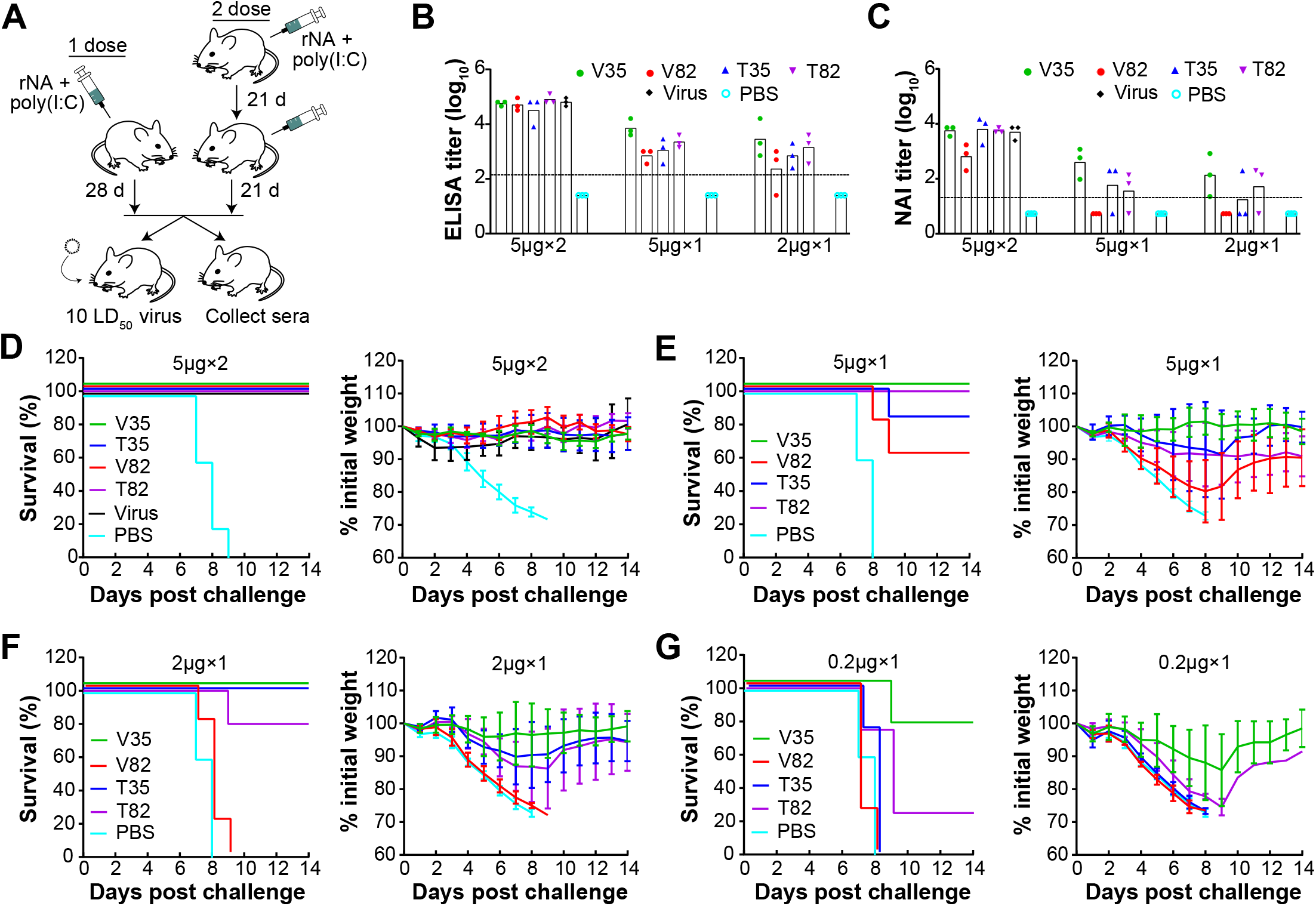
N1-BR18 rNAs elicit different protective immune responses in mice. **(A)** Mice received two i.m. injections 21-days apart with 5 μg rNA or a single-dose i.m. immunization with 5, 2, or 0.2 μg rNA. On day 21 after the second dose or day 28 after the single-dose immunization, 3 mice from each group (except those immunized with 0.2 μg rNA) were euthanized to collect sera for antibody assessment, while the remaining mice were challenged with 10 LD_50_ of H6N1_BR18_×PR8 virus and monitored for weight loss and mortality for up to 14 days. Purified BR18×WSN virus was used as a positive control in the two-dose immunizations, PBS was used as the negative control, and all immunizations were conducted with 5 μg poly(I:C) adjuvant. **(B)** Serum NA-binding antibody titers measured with ELISA and **(C)** Serum NAI antibody titers measured with ELLA. H6N1_BR18_×PR8 virus was used as the antigen for both the ELISA and ELLA measurements. The limits of detection (denoted by the dotted lines) for ELISA and ELLA were 125 (2.1 log_10_) and 20 (1.3 log_10_), respectively, and titers below these limits were arbitrarily set to 25 (1.4 log_10_) for ELISA and 5 (0.7 log_10_) for ELLA for the purpose of calculation. Shown are the mean ± SD (n=3) of data from two technical repeats. **(D)** Survival (left panel) and weight loss (right panel) of mice (n=5) challenged with virus after two-dose immunizations with 5 μg of the indicated rNA. **(E-G)** Survival and weight loss of mice challenged with virus after a single immunization with (**E**) 5 μg, (**F**) 2 μg, or (**G**) 0.2 μg of the indicated rNA. (n=5 in the 5 and 2 μg groups, n= 4 in the 0.2 μg group).

Based on the robust antibody responses from the single-dose immunizations, we also included groups of mice immunized with 0.2 μg of rNA adjuvanted with poly(I:C) in the lethal viral challenge experiment. In the groups that received two 5 μg doses of rNA or purified BR18×WSN virus, all mice were protected and displayed no evident weight loss, whereas the control group immunized with PBS containing poly(I:C) succumbed to the viral challenge (Fig. 5D). In the single-dose groups, all mice immunized with 5 or 2 μg of V35 survived the viral challenge with little weight loss, and most mice that received 0.2 μg of V35 also survived with a maximal average weight loss of ~15% (Fig. 5E-G). T35 and T82 protected almost all the animals at a single dose of 5 and 2 μg, although substantial weight loss was observed in these groups, and the 0.2 μg dose showed little or no protection (Fig. 5E-G). Consistent with the poor antibody response, V82 provided the least protection across all of the single-dose immunizations. Together, these results show that the domains included in the rNA design are crucial for achieving optimal protection, and the unexpected difference in protection from V82 and V35 indicate that enzymatic activity is not necessarily predictive of protection of rNA.

## DISCUSSION

Since the 1990s, the biochemical and immunological properties of rNA have been studied in some detail (19, 21, 30, 47). These and more recent studies (20, 28, 29, 48) have established that rNA can be protective and provide cross protection against influenza strains carrying a similar NA but different HA subtypes, leading to the proposal that rNA could be used to supplement existing influenza vaccines. However, all the previously reported animal studies used a two-dose (prime and boost) immunization regimen with high rNA amounts and provided little information on the rNA quality attributes, which are critical for evaluating a vaccine antigen. The results from our systematic comparison demonstrate the following: *-i-* Enzymatic properties of rNAs are dependent on the NA domains included in the construct design; *-ii-* Molecular size stability of the rNA is influenced by the properties of the tetramerization domain; *-iii-* Single-dose rNA immunizations in mice can provide full protection against a lethal viral challenge and are more informative for evaluating rNAs; and *-iv-* Protective efficacy can substantially differ between rNA designs with similar attributes, indicating that the rNA immunogenicity is mainly determined by the combination of the NA and the tetramerization domains. These findings show that the rNA design is critical for optimal protective efficacy and that rNA antigens being developed to improve influenza vaccines would benefit from a comprehensive evaluation.

The interesting observation that rNA designs including the head-domain (V82 and T82) generate a higher percentage of functional rNA than designs containing the full-ectodomain (V35 and T35) suggests that the stalk region impairs the function of the head-domain. Earlier studies also reported that the presence of the stalk significantly reduces the activity of a secreted NA without a tetramerization domain, implying that head-domain assembly is most efficient when the stalk is attached to a lipid bilayer by the tetrameric amphipathic transmembrane (49, 50). To complement the absence of the transmembrane domain, the four rNAs in this study contain an *N*-terminal tetramerization domain and they are all recognized similarly by various N1 head-specific mAbs, indicating multiple epitopes in the head-domain are largely preserved despite the activity differences. In addition, the rNA designs (V35 and T35) that produce the lowest percentage of functional NA elicited comparable (T35 versus T82) or higher antibody response and protection (V35 versus V82). All these results raise the question of what causes the decrease in enzymatic activity of the rNAs containing the full-ectodomain. Previous work has shown that full-length NA assembles through a cooperative process that requires compatibility between the head and transmembrane domain and that formation of tetramer-dependent central Ca^2+^ binding pocket is essential for NA activity (38, 51). We speculate that the tetramerization domains attached to the stalk are less compatible with the head domain than the native transmembrane region, resulting in the suboptimal formation of the central Ca^2+^ binding pocket and hence the lower activity.

We also observed that the tetramerization domain from VASP introduces more instability in the molecular size of the rNAs than the one from TB. This phenotype was especially evident during storage at 4°C where V35 and V82 showed time-dependent shifts in molecular size, which is indicative of higher order oligomer or multimer formation. The molecular size increase did not coincide with a loss in activity (data not shown), suggesting that the number of functional active sites in the higher order V35 and V82 oligomers did not change. The more stable T35 and T82 might be preferred from the perspective of vaccine manufacture as most influenza vaccines are commonly stored at 4°C. However, it is unknown if the increase in molecular size, as observed for V35 and V82 stored at 4°C, is a beneficial attribute for immunogenicity since our protection experiments were performed with rNAs stored at −80°C prior to the shift in size. Protection experiments aimed at assessing the impact of the molecular size increase on the immune response and protection will be reported in a subsequent study, which will help to determine the rNA storage requirements

With the common two-dose approach and high amounts of rNA the protection in mice were almost indiscernible, but with the single-dose immunizations clear differences were observed in the protection from the rNAs, suggesting that multiple doses of high amounts may mask difference in the quality of the rNA antigens. Following the single-dose immunizations, the highest antibody titers and protective efficacy were observed for V35, which produces a low percentage of enzymatically active NA, whereas the poorest immune response was observed for V82 that produces a high percentage of enzymatically active NA. In contrast, similar antibody titers and protective efficacy were observed for T35 and T82. The differences in the protective efficacy between V82 and the other three rNAs, especially T82, were very unexpected. We did observe a higher prevalence of a particular glycoform on V82 compared to T82. However, it is unlikely that this minor modification of an insect cell glycoform is solely responsible for the low immunogenicity of V82. The probable factors could be the smaller molecular size of V82, the propensity for V82 to form multimers, and the immunogenicity of the charged VASP domain in the context of the smaller NA-head domain. Our results emphasize that enzymatically active rNAs are not necessarily the most immunogenic, which is significantly different than the common belief that enzymatic activity is an ideal attribute for assessing NA quality.

In summary, our findings highlight the necessity to carefully select the elements included in the design of rNAs, as different attributes are influenced by the choice of the tetramerization and NA domains. Our data also indicate that NA activity may not be the best attribute for assessing the quality of rNAs. Based on the clear difference across the four recombinant N1 constructs, future studies are needed to evaluate the designs of other vaccine relevant rNAs, to identify additional methods for optimizing rNA production and to assess whether supplementing influenza vaccines with rNA can enhance the immunogenicity and reduce the HA dose amounts needed in the vaccine.

## MATERIALS AND METHODS

### Cells and viruses

High Five insect cells (Invitrogen) maintained in Express Five serum free medium (Life Technologies) were used for the N1-BR18 rNA expression. Recombinant baculoviruses that express the N1-BR18 rNAs (see Fig. 1B for the rNA construct design) were produced by GenScript Inc. Recombinant influenza viruses were rescued in Madin-Darby canine kidney cells and human embryonic kidney 293T cells using reverse genetics as previously reported (52, 53). These viruses included BR18×WSN, which bears the HA and NA genes from BR18 (H1N1) and the internal genes from A/WSN/1933 (WSN, H1N1), H6N1_BR18_×WSN and H6N1_BR18_×PR8, which bear the HA gene from A/turkey/Massachusetts/3740/1965 (H6N2), the NA gene from BR18, and the internal genes from WSN and A/Puerto Rico/8/1934 (PR8, H1N1), respectively. The rescued viruses were propagated in 9-11-day-old specific pathogen-free embryonated chicken eggs. The median lethal dose (LD_50_) of H6N1_BR18_ in mice was determined for the lethal viral challenge. Viruses were also inactivated with *β*-propiolactone (Sigma) and purified by sucrose gradient centrifugation for *in vitro* assays and the animal study.

### rNA expression and purification

High Five insect cells were grown to a density of ~2 ×10^6^ cell/ml in shaker flasks at 120 rpm and 27.5°C prior to infection with each recombinant baculovirus at a multiplicity of infection of ~2.0-5.0. At 72-96 h post-infection, when the NA activity plateaued, the cell culture supernatant was clarified, concentrated and exchanged to a pH 8.0 buffer (50 mM Tris, 300 mM NaCl, 1 mM CaCl_2_) by tangential flow filtration using a cartridge with a 30-kD molecular weight cutoff. The buffer was then adjusted to contain 40 mM imidazole and the rNAs were purified with a HisTrap^TM^ FF 1 ml column (GE Healthcare) using an Akta Start protein purification system (Cytiva). Alternatively, rNAs were purified using a HisTrap^TM^ FF 5 ml column (GE Healthcare) from the cell culture supernatant that was clarified but not concentrated or buffer exchanged. The column was washed with a pH 8.0 buffer (50 mM Tris, 300 mM NaCl, 1 mM CaCl_2_) containing 40 mM imidazole and the bound protein was eluted using a pH 8.0 buffer (50 mM Tris, 300 mM NaCl, 1 mM CaCl_2_) supplemented with 250 mM imidazole. Fractions containing the rNAs were pooled and exchanged into in a pH 6.5 buffer (50 mM Tris, 150 mM NaCl, 1 mM CaCl_2_, 5% glycerol) using 15ml centrifugal filters with a 30-kD molecular weight cutoff (Millipore) and the rNA concentration was measured and adjusted to ~1.0 mg/ml. Each rNA was then aliquoted and stored at −80°C or 4°C for subsequent assays.

### SDS-PAGE and Western blot

rNAs (2 μg/lane for Coomassie, 0.2 μg/lane for Western blot) were mixed with 2× sample buffer containing 50 mM DTT, heated at 50°C for 10 min, and resolved by 4-12 % polyacrylamide Tris-Glycine SDS-PAGE wedge gels (Thermo Fisher Scientific). Gels were either stained with simple blue (Thermo Fisher Scientific) or transferred to a 0.45-μm pore PVDF membrane (Life Technologies) at 65 V for 1h. The membrane was blocked with the AzureSpectra Fluorescence Blot Blocking Buffer (Azure Biosystems), and incubated with 1 μg/ml N1-specific rabbit mAb (Sino Biological) and AzureSpectra 700 goat-anti-rabbit IgG (Azure Biosystems). The Coomassie gels and immunoblots were then imaged using an Azure C600 Bioanalytical Imaging System (Azure Biosystems).

### Glycan analysis

Each rNA was exchanged into 50 mM ammonium bicarbonate buffer, pH 8.0, using 0.5 ml centrifugal filters (Millipore). The rNA samples were reduced by the addition of 5 mM DTT followed by a 30 min incubation at 60°C. The cysteines were then alkylated by incubation with 15 mM iodoacetamide for 30 min at room temperature in the dark. The alkylation reactions were quenched by adding 25 mM DTT, and 25 μg of each rNA sample was digested at 37°C overnight with trypsin (Promega). After 10 min at 95°C to denature the trypsin, the samples were incubated at 37°C overnight with PNGase F (New England BioLabs) to release the glycans. Glycan purification, permethylation, data collection and analysis were done as described previously (54), and the glycoform assignments were determined using a reference library of glycans known to be present in insect cells.

### ELISA

A sandwich ELISA was performed as previously described (41) to confirm the presence of various epitopes on N1-BR18 rNAs. Briefly, N1-specific mAbs CD6, 4C4, 1H5 (42–44) and an N2-specific mAb B10 (38) were coated onto Immulon^®^ 2HB flat bottom microtiter plates (Thermo Fisher Scientific) at 1 μg/well. After blocking with 15% fetal bovine serum (FBS) (Atlanta Biologics) in PBS, diluted N1-BR18 rNAs and BR18×WSN virus were added and incubated at 37°C for 1 h, followed by washing and incubation with HRP-conjugated mAb 4E9 or 4C4 at 37°C for 1 h. Plates were then developed using the substrate *o*-phenylenediamine dihydrochloride (OPD; Sigma) for 10 min, the reactions were stopped with 1 N H_2_SO_4_, and the absorbance values at 490 nm (Abs_490_) were read. To measure the NA-binding antibody titers in mouse serum samples, purified H6N1_BR18_×PR8 virus was coated onto plates at 0.5 μg/well of the total viral protein. After blocking with 15% FBS in PBS, 2-fold serially diluted serum samples were added and incubated at 37°C for 1 h. The plates were then washed, HRP-conjugated goat-anti-mouse IgG (Sigma) was added, and the plates were incubated at 37°C for 1h. The plates were developed the same way as for the sandwich ELISA., The cutoff Abs_490_ value was set at 0.08.

### Enzymatic activity assay

To examine the NA activity, rNAs and BR18×WSN virus were diluted in 25 μl in 96-well, black wall, clear bottom plates (Corning) and warmed to 37°C. The reaction was initiated by mixing each sample with 175 μl of substrate solution [170 μl 0.1 M KH_2_PO_4_ containing 1 mM CaCl_2_ (pH 6.0) and 5 μl of 2 mM Mu-NANA]. The fluorescence was measured using a Cytation 5 Cell Imaging Multi-Mode Reader (Biotek) at 37°C for 10 min using 30 sec intervals and a 365 nm excitation wavelength and a 450 nm emission wavelength. The NA activity was determined based on the slope of the early linear region in the emission versus time graph.

The Michaelis-Menten kinetic analysis was performed by diluting the rNAs and BR18×WSN virus to a known concentration and measuring the enzymatic activity with the presence of increasing concentrations of Mu-NANA the reached saturation. The *V*_max_ and *K*_m_ values were calculated by analyzing the nonlinear fitting curves with GraphPad Prism version 8.0 (GraphPad Software). The *k*_cat_ values were calculated using the *V*_max_ value, the rNA concentration and a 4-methylumbelliferone (Sigma) standard curve.

### TR1 assay

The TR1 assay was performed as previously reported (40) with modifications. In brief, F_2_Mu was 2-fold serially diluted in 96-well, black wall, clear bottom plates (Corning), starting from 1.0 μM, in 200 μl pH 7.6 buffer (50 mM Tris, 20 mM CaCl_2_). The fluorescence signals were read at a 365 nm excitation wavelength and a 450 nm emission wavelength using a Cytation 5 Cell Imaging Multi-Mode Reader (Biotek) and a standard curve was created. The rNAs and were serially diluted to 20 μl using pH 7.6 buffer (50 mM Tris, 20 mM CaCl_2_) in 96-well, black wall, clear bottom plates (Corning), and the reactions were initiated by adding 180 μl TR1 solution (175 μl 50 mM Tris pH 7.6 containing 20 mM CaCl_2_, 5 μl 1 mM TR1). The fluorescence signals were monitored continuously for 30 min at 30 sec intervals, and the number of the active NA catalytic sites was calculated based on the signals reached at the plateau and the standard calibration equation.

### rNA stability analysis

The thermostability was monitored by incubating the rNAs at 37, 43.2, 52, 54.1, 56.4, 59.6, 63.7, 67.1 and 70°C for 10 min in a C1000 Touch^TM^ Cycler (Bio-Rad) and measuring the enzymatic activity with the Mu-NANA assay. The data were then analyzed using nonlinear fitting curves to calculate the *T_50_* values on GraphPad Prism version 8.0 (GraphPad Software). The freeze-thaw stability was determined by measuring the enzymatic activity of rNAs following 5 cycles of freezing for 10 min on dry ice and thawing briefly at 37°C, the enzymatic activity of the treated samples was normalized to that of untreated samples. To examine the molecular size stability at the routine storage temperature, rNAs stored at −80°C were thawed at 1, 3, and 6-months and analyzed by SEC. Briefly, 10 μl of each rNA, adjusted to 0.5 mg/ml with the pH 6.5 buffer (50 mM Tris, 150 mM NaCl, 1 mM CaCl_2_, 5% glycerol) was analyzed using an Agilent 1260 prime HPLC equipped with an AdvanceBio SEC 300Å column, a variable wavelength detector set at 220 and 280 nm, and a fraction collector, run at a flow rate of 1 ml/min. The molecular weights for each rNA were estimated using an AdvanceBio SEC 300Å protein standard (Agilent) of known molecular weights that was included in the run and the presence of NA in each fraction was measured with the Mu-NANA assay.

### Animal study

Mouse experiments were conducted to examine the immunogenicity and protective efficacy of N1-BR18 rNAs against lethal viral challenge. For the two-dose immunization regimen, DBA/2 mice (female, 6-wk old; The Jackson Laboratory; n=8 per group) were immunized i.m. with each rNA 5 μg mixed with 5 μg poly(I:C) adjuvant (Sigma) and boosted with the same dose of rNA and poly(I:C) at a 21-day interval. On day 21 post-boost, 3 mice from each group were euthanized, and the blood was collected for measuring the NA-binding antibodies with ELISA and NAI antibodies with ELLA. The remaining 5 mice per group were challenged intranasally (i.n.) with 10 LD_50_ of H6N1_BR18_×PR8 in 50 μl of PBS. These mice were monitored for weight loss and mortality for up to 14 days, and mice that lost 25% weight were euthanized. Mice primed and boosted with 5 μg inactivated, purified BR18×WSN virus adjuvanted with 5 μg poly(I:C) were included as the positive control and mice receiving PBS containing 5 μg poly(I:C) were included as the negative control. For the single-dose regimen, DBA/2 mice (female, 8-wk old; The Jackson Laboratory; n=8 for the 5 and 2 μg rNA groups, n=4 for the 0.2 μg rNA groups) received a single i.m. immunization of rNAs at 5, 2, 0.2 μg, or PBS mixed with 5 μg poly(I:C). On day 28 post-immunization, 3 mice from each group immunized with 5 or 2 μg rNA were euthanized for blood collection and the NA serum antibody titers were measured. The remaining animals (n=5 or 4) in each group were challenged and monitored similarly. Federal guidelines and protocols approved by the Food and Drug Administration Institutional Animal Care and Use Committee were followed in the animal experiments.

## NAI assay

The NAI antibody titers in mouse serum samples were measured with ELLA as described previously (55). Serial dilutions of the serum samples were mixed with a predetermined amount of virus diluted in pH 6.5 MES buffer (KD Medical) containing 1% bovine serum albumin (Sigma) and 0.5% Tween-20 (Sigma). The mixture was added to 96-well plates (Thermo Fisher Scientific) coated with 2.5 μg/well of fetuin (Sigma) and incubated overnight at 37°C. Plates were washed with PBS containing 0.05% Tween-20 (PBST), followed by adding HRP-conjugated peanut agglutinin (Sigma). Plates were incubated at room temperature for 2 h in the dark and washed with PBST before the addition of the OPD substrate. The reaction was stopped by adding 1 N H_2_SO_4_ and Abs_490_ values were read, the antibody titer was expressed as the reciprocal of the highest dilution that exhibited ≥ 50% inhibition of NA activity.

## ACKOWLEDGEMENTS

This work was supported by intramural funds from the Food and Drug Administration. We thank St. Jude Children’s Research Hospital for providing plasmids that were used to rescue viruses. We thank Paul Carney and James Stevens from the Centers for Disease Control and Prevention for technical help and providing the N2 protein used in the study. We are indebted to staff of the Division of Veterinary Services, Center for Biologics Evaluation and Research, Food and Drug Administration, for excellent animal care. The findings and conclusions in this report are those of the authors and do not necessarily represent the views of the Food and Drug Administration.

R.D. and H.W. designed the study. J.G., L.K., L.P., T.M., J.P., Z.G. and H.W. performed and/or helped with the experiments. R.D. and H.W. wrote the paper. S.G.W., J.C., R.D. and H.W. edited the paper.

Disclosures: S.G.W. and Z.G. are named contributors to a patent application submitted by the University of British Columbia concerning the development and use of TR1 as an active site titrant for influenza NA. The authors have no additional financial interests.

## REFERENCES

1. Wu NC, Zost SJ, Thompson AJ, Oyen D, Nycholat CM, McBride R, Paulson JC, Hensley SE, Wilson IA. 2017. A structural explanation for the low effectiveness of the seasonal influenza H3N2 vaccine. PLoS Pathog 13:e1006682.

2. Paules CI, Sullivan SG, Subbarao K, Fauci AS. 2018. Chasing Seasonal Influenza - The Need for a Universal Influenza Vaccine. N Engl J Med 378:7–9.

3. Zost SJ, Parkhouse K, Gumina ME, Kim K, Diaz Perez S, Wilson PC, Treanor JJ, Sant AJ, Cobey S, Hensley SE. 2017. Contemporary H3N2 influenza viruses have a glycosylation site that alters binding of antibodies elicited by egg-adapted vaccine strains. Proc Natl Acad Sci U S A 114:12578–12583.

4. Ostbye H, Gao J, Martinez MR, Wang H, de Gier JW, Daniels R. 2020. N-Linked Glycan Sites on the Influenza A Virus Neuraminidase Head Domain Are Required for Efficient Viral Incorporation and Replication. J Virol 94.

5. Gaymard A, Le Briand N, Frobert E, Lina B, Escuret V. 2016. Functional balance between neuraminidase and haemagglutinin in influenza viruses. Clin Microbiol Infect 22:975–983.

6. Dou D, Revol R, Ostbye H, Wang H, Daniels R. 2018. Influenza A Virus Cell Entry, Replication, Virion Assembly and Movement. Front Immunol 9:1581.

7. Murphy BR, Chalhub EG, Nusinoff SR, Chanock RM. 1972. Temperature-sensitive mutants of influenza virus. II. Attenuation of ts recombinants for man. J Infect Dis 126:170–8.

8. Schulman JL, Khakpour M, Kilbourne ED. 1968. Protective effects of specific immunity to viral neuraminidase on influenza virus infection of mice. J Virol 2:778–86.

9. Allan WH, Madeley CR, Kendal AP. 1971. Studies with avian influenza A viruses: cross protection experiments in chickens. J Gen Virol 12:79–84.

10. Monto AS, Kendal AP. 1973. Effect of neuraminidase antibody on Hong Kong influenza. Lancet 1:623–5.

11. Couch RB, Kasel JA, Gerin JL, Schulman JL, Kilbourne ED. 1974. Induction of partial immunity to influenza by a neuraminidase-specific vaccine. J Infect Dis 129:411–20.

12. Memoli MJ, Shaw PA, Han A, Czajkowski L, Reed S, Athota R, Bristol T, Fargis S, Risos K, Powers JH, Davey RT, Jr., Taubenberger JK. 2016. Evaluation of Antihemagglutinin and Antineuraminidase Antibodies as Correlates of Protection in an Influenza A/H1N1 Virus Healthy Human Challenge Model. mBio 7:e00417–16.

13. Air GM. 2012. Influenza neuraminidase. Influenza Other Respi Viruses 6:245–56.

14. Kilbourne ED, Johansson BE, Grajower B. 1990. Independent and disparate evolution in nature of influenza A virus hemagglutinin and neuraminidase glycoproteins. Proc Natl Acad Sci U S A 87:786–90.

15. Krammer F, Fouchier RAM, Eichelberger MC, Webby RJ, Shaw-Saliba K, Wan H, Wilson PC, Compans RW, Skountzou I, Monto AS. 2018. NAction! How Can Neuraminidase-Based Immunity Contribute to Better Influenza Virus Vaccines? mBio 9.

16. Eichelberger MC, Monto AS. 2019. Neuraminidase, the Forgotten Surface Antigen, Emerges as an Influenza Vaccine Target for Broadened Protection. J Infect Dis 219:S75–S80.

17. Wang H, Dou D, Ostbye H, Revol R, Daniels R. 2019. Structural restrictions for influenza neuraminidase activity promote adaptation and diversification. Nat Microbiol 4:2565–2577.

18. Brett IC, Johansson BE. 2005. Immunization against influenza A virus: comparison of conventional inactivated, live-attenuated and recombinant baculovirus produced purified hemagglutinin and neuraminidase vaccines in a murine model system. Virology 339:273–80.

19. Deroo T, Jou WM, Fiers W. 1996. Recombinant neuraminidase vaccine protects against lethal influenza. Vaccine 14:561–9.

20. Job ER, Ysenbaert T, Smet A, Christopoulou I, Strugnell T, Oloo EO, Oomen RP, Kleanthous H, Vogel TU, Saelens X. 2018. Broadened immunity against influenza by vaccination with computationally designed influenza virus N1 neuraminidase constructs. NPJ Vaccines 3:55.

21. Martinet W, Saelens X, Deroo T, Neirynck S, Contreras R, Min Jou W, Fiers W. 1997. Protection of mice against a lethal influenza challenge by immunization with yeast-derived recombinant influenza neuraminidase. Eur J Biochem 247:332–8.

22. Johansson BE, Matthews JT, Kilbourne ED. 1998. Supplementation of conventional influenza A vaccine with purified viral neuraminidase results in a balanced and broadened immune response. Vaccine 16:1009–15.

23. Johansson BE, Bucher DJ, Kilbourne ED. 1989. Purified influenza virus hemagglutinin and neuraminidase are equivalent in stimulation of antibody response but induce contrasting types of immunity to infection. J Virol 63:1239–46.

24. Kilbourne ED, Couch RB, Kasel JA, Keitel WA, Cate TR, Quarles JH, Grajower B, Pokorny BA, Johansson BE. 1995. Purified influenza A virus N2 neuraminidase vaccine is immunogenic and non-toxic in humans. Vaccine 13:1799–803.

25. Izikson R, Leffell DJ, Bock SA, Patriarca PA, Post P, Dunkle LM, Cox MM. 2015. Randomized comparison of the safety of Flublok((R)) versus licensed inactivated influenza vaccine in healthy, medically stable adults >= 50 years of age. Vaccine 33:6622–8.

26. Johansson BE, Grajower B, Kilbourne ED. 1993. Infection-permissive immunization with influenza virus neuraminidase prevents weight loss in infected mice. Vaccine 11:1037–9.

27. Johansson BE, Pokorny BA, Tiso VA. 2002. Supplementation of conventional trivalent influenza vaccine with purified viral N1 and N2 neuraminidases induces a balanced immune response without antigenic competition. Vaccine 20:1670–4.

28. Wohlbold TJ, Nachbagauer R, Xu H, Tan GS, Hirsh A, Brokstad KA, Cox RJ, Palese P, Krammer F. 2015. Vaccination with adjuvanted recombinant neuraminidase induces broad heterologous, but not heterosubtypic, cross-protection against influenza virus infection in mice. mBio 6:e02556.

29. McMahon M, Strohmeier S, Rajendran M, Capuano C, Ellebedy AH, Wilson PC, Krammer F. 2020. Correctly folded - but not necessarily functional - influenza virus neuraminidase is required to induce protective antibody responses in mice. Vaccine 38:7129–7137.

30. Johansson BE, Price PM, Kilbourne ED. 1995. Immunogenicity of influenza A virus N2 neuraminidase produced in insect larvae by baculovirus recombinants. Vaccine 13:841–5.

31. Varghese JN, Laver WG, Colman PM. 1983. Structure of the influenza virus glycoprotein antigen neuraminidase at 2.9 A resolution. Nature 303:35–40.

32. Paterson RG, Lamb RA. 1990. Conversion of a class II integral membrane protein into a soluble and efficiently secreted protein: multiple intracellular and extracellular oligomeric and conformational forms. J Cell Biol 110:999–1011.

33. Xu X, Zhu X, Dwek RA, Stevens J, Wilson IA. 2008. Structural characterization of the 1918 influenza virus H1N1 neuraminidase. J Virol 82:10493–501.

34. Margine I, Palese P, Krammer F. 2013. Expression of functional recombinant hemagglutinin and neuraminidase proteins from the novel H7N9 influenza virus using the baculovirus expression system. J Vis Exp doi:10.3791/51112:e51112.

35. Johansson BE. 1999. Immunization with influenza A virus hemagglutinin and neuraminidase produced in recombinant baculovirus results in a balanced and broadened immune response superior to conventional vaccine. Vaccine 17:2073–80.

36. Dai M, Guo H, Dortmans JC, Dekkers J, Nordholm J, Daniels R, van Kuppeveld FJ, de Vries E, de Haan CA. 2016. Identification of Residues That Affect Oligomerization and/or Enzymatic Activity of Influenza Virus H5N1 Neuraminidase Proteins. J Virol 90:9457–70.

37. Wang N, Glidden EJ, Murphy SR, Pearse BR, Hebert DN. 2008. The cotranslational maturation program for the type II membrane glycoprotein influenza neuraminidase. J Biol Chem 283:33826–37.

38. Wan H, Gao J, Yang H, Yang S, Harvey R, Chen YQ, Zheng NY, Chang J, Carney PJ, Li X, Plant E, Jiang L, Couzens L, Wang C, Strohmeier S, Wu WW, Shen RF, Krammer F, Cipollo JF, Wilson PC, Stevens J, Wan XF, Eichelberger MC, Ye Z. 2019. The neuraminidase of A(H3N2) influenza viruses circulating since 2016 is antigenically distinct from the A/Hong Kong/4801/2014 vaccine strain. Nat Microbiol 4:2216–2225.

39. Gao Z, Niikura M, Withers SG. 2017. Ultrasensitive Fluorogenic Reagents for Neuraminidase Titration. Angew Chem Int Ed Engl 56:6112–6116.

40. Gao Z, Robinson K, Skowronski DM, De Serres G, Withers SG. 2020. Quantification of the total neuraminidase content of recent commercially-available influenza vaccines: Introducing a neuraminidase titration reagent. Vaccine 38:715–718.

41. Wan H, Sultana I, Couzens LK, Mindaye S, Eichelberger MC. 2017. Assessment of influenza A neuraminidase (subtype N1) potency by ELISA. J Virol Methods 244:23–28.

42. Wan H, Yang H, Shore DA, Garten RJ, Couzens L, Gao J, Jiang L, Carney PJ, Villanueva J, Stevens J, Eichelberger MC. 2015. Structural characterization of a protective epitope spanning A(H1N1)pdm09 influenza virus neuraminidase monomers. Nat Commun 6:6114.

43. Wan H, Gao J, Xu K, Chen H, Couzens LK, Rivers KH, Easterbrook JD, Yang K, Zhong L, Rajabi M, Ye J, Sultana I, Wan XF, Liu X, Perez DR, Taubenberger JK, Eichelberger MC. 2013. Molecular basis for broad neuraminidase immunity: conserved epitopes in seasonal and pandemic H1N1 as well as H5N1 influenza viruses. J Virol 87:9290–300.

44. Jiang L, Fantoni G, Couzens L, Gao J, Plant E, Ye Z, Eichelberger MC, Wan H. 2016. Comparative Efficacy of Monoclonal Antibodies That Bind to Different Epitopes of the 2009 Pandemic H1N1 Influenza Virus Neuraminidase. J Virol 90:117–28.

45. Kilbourne ED, Pokorny BA, Johansson B, Brett I, Milev Y, Matthews JT. 2004. Protection of mice with recombinant influenza virus neuraminidase. J Infect Dis 189:459–61.

46. Bosch BJ, Bodewes R, de Vries RP, Kreijtz JH, Bartelink W, van Amerongen G, Rimmelzwaan GF, de Haan CA, Osterhaus AD, Rottier PJ. 2010. Recombinant soluble, multimeric HA and NA exhibit distinctive types of protection against pandemic swine-origin 2009 A(H1N1) influenza virus infection in ferrets. J Virol 84:10366–74.

47. Mather KA, White JF, Hudson PJ, McKimm-Breschkin JL. 1992. Expression of influenza neuraminidase in baculovirus-infected cells. Virus Res 26:127–39.

48. Giurgea LT, Park JK, Walters KA, Scherler K, Cervantes-Medina A, Freeman A, Rosas LA, Kash JC, Taubenberger JK, Memoli MJ. 2021. The effect of calcium and magnesium on activity, immunogenicity, and efficacy of a recombinant N1/N2 neuraminidase vaccine. NPJ Vaccines 6:48.

49. da Silva DV, Nordholm J, Madjo U, Pfeiffer A, Daniels R. 2013. Assembly of subtype 1 influenza neuraminidase is driven by both the transmembrane and head domains. J Biol Chem 288:644–53.

50. Nordholm J, da Silva DV, Damjanovic J, Dou D, Daniels R. 2013. Polar residues and their positional context dictate the transmembrane domain interactions of influenza A neuraminidases. J Biol Chem 288:10652–60.

51. da Silva DV, Nordholm J, Dou D, Wang H, Rossman JS, Daniels R. 2015. The influenza virus neuraminidase protein transmembrane and head domains have coevolved. J Virol 89:1094–104.

52. Neumann G, Watanabe T, Ito H, Watanabe S, Goto H, Gao P, Hughes M, Perez DR, Donis R, Hoffmann E, Hobom G, Kawaoka Y. 1999. Generation of influenza A viruses entirely from cloned cDNAs. Proc Natl Acad Sci U S A 96:9345–50.

53. Sandbulte MR, Gao J, Straight TM, Eichelberger MC. 2009. A miniaturized assay for influenza neuraminidase-inhibiting antibodies utilizing reverse genetics-derived antigens. Influenza Other Respir Viruses 3:233–40.

54. Parsons LM, Bouwman KM, Azurmendi H, de Vries RP, Cipollo JF, Verheije MH. 2019. Glycosylation of the viral attachment protein of avian coronavirus is essential for host cell and receptor binding. J Biol Chem 294:7797–7809.

55. Gao J, Couzens L, Eichelberger MC. 2016. Measuring Influenza Neuraminidase Inhibition Antibody Titers by Enzyme-linked Lectin Assay. J Vis Exp doi:10.3791/54573.

